# The duration of *Eragrostis capillaris* is not strictly annual: it does not senesce, can reproduce multiple times, but lacks frost tolerance

**DOI:** 10.1101/2022.06.02.494534

**Authors:** Gordon G. McNickle

**Affiliations:** Department of Botany and Plant Pathology, Purdue University, West Lafayette, IN 47907-2054; Center for Plant Biology, Purdue University, West Lafayette, IN 47907-2054

## Abstract

The duration of *Eragrostis capillaris* (L.) Nees (love lacegrass) is currently described as an annual by all sources. Here, I describe observations that show it does not senesce naturally, can reproduce multiple times across multiple years, but lacks frost tolerance. I conclude that *E. capillaris* probably grows as an annual throughout much of its range due to killing frost, but can have a perennial duration in locations or years that do not have a killing frost. Its status as an annual is likely to change under a warming climate. I suggest its duration should be updated to “annual or perennial” in taxonomic guides. It remains an open question how common such shifts might be under a warming climate.

## INTRODUCTION

There are many strictly annual plants that germinate, reproduce and then senesce in one growing season. Some annuals (E.g. the model plant *Arabidopsis thaliana* L.) complete their life cycle remarkably rapidly germinating and then naturally senescing in just a fraction of the total growing season (Koornneef & Meinke 2010). Alternatively, many long-lived perennials do not reproduce in their first year(s), and instead focus on establishing vegetative organs such as roots or secondary stem growth in early years prior to allocating to reproduction (Friedman & Rubin 2015). However, for some species the line between annual and perennial is not discrete.

Some perennials are capable of germinating and reproducing in one growing season, and some annuals can live much longer than one year in greenhouse conditions (Fox 1990). In these cases, the line between annual and perennial duration blurs depending on causes of mortality in nature. In these blurred cases, the duration of such plants is frequently described as “annual or perennial” (E.g. Usda & Nrcs 2022). Most often, a lack of environmental tolerance (E.g. frost, drought, flooding) leads to a potentially perennial plant that can reproduce within a single growing season having an annual duration in some locations or years (Beatley 1970; Sano, Morishima & Oka 1980; Fox 1990).

Every source of which I am aware lists the duration of *Eragrostis capillaris* (L.) Nees (love lacegrass) as purely annual (E.g. Bourfford 1993+; Usda & Nrcs 2022). However, after working with it for some greenhouse experiments, I observed that *E. capillaris* appears to have no natural mechanism for senescence. I hypothesized that it lacks frost tolerance which might allow it to grow as either an annual or a perennial in some locations or years that lack a killing frost.

## METHODS

In March of 2021 I started 12 pots with *E. capillaris* seeds sourced from Roundstone Native Seed Company (Upton, KY, USA). Pots were 9.5 litres (23 cm diameter, 23 cm tall), the growing medium was 1:1 sand and potting soil (propagation mix soil, Sungro Company, Agawam, Massachusetts, USA). One tablespoon of slow release fertilizer with micronutrients (Osmocote plus, The Scotts Company L.L.C., Marysville, OH, USA) was added to each pot in April of 2021 after plants had established. Pots were watered freely twice per week; no supplemental lighting was used. By November 2021 (9 months of growth) there was no evidence of senescence (Fig 1A). In my experience clipping annual grasses has more severe consequences than clipping perennials. Thus, all plants were clipped down to 1 cm height (Fig 1B) and the following treatments were imposed to investigate frost tolerance and senescence:

**FIGURE 1:**
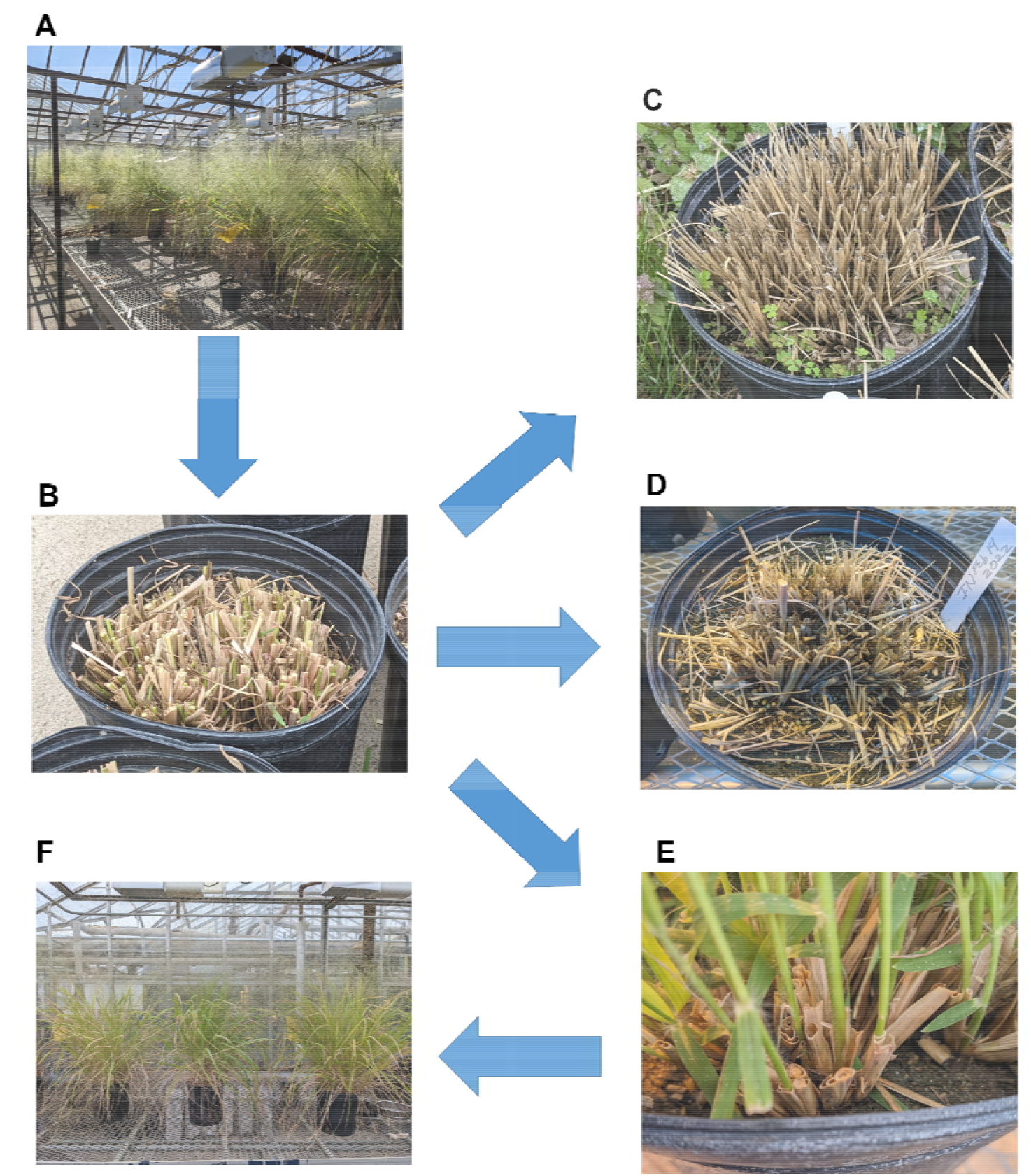
Photographs of the different experimental manipulations performed in this study. Arrows represent time; see main text for timings of each experiment. (A) Originally plants grew from March to November in 2021. (B) In November 2021 all plants were clipped to 1cm. (C) For the plants that were placed outside and experienced a natural winter, there was no regrowth by May of 2022. (D) For the plants that experienced either four or eight weeks in a −20°C freezer, there was no regrowth after eight weeks in the greenhouse following removal from the freezer. (E) Plants that were clipped and placed back in the greenhouse re-sprouted from clipped stems, and (F) by April 2022 (14 months after germination) the had flowered a second time.

(i) Six pots were placed outside (West Lafayette, Indiana, USA, 40°42’26.0”N, 86°91’88.2”W) and kept moist all winter; (ii) three pots were placed in a −20°C freezer for four weeks and then returned to the greenhouse; (iii) three pots were placed in a −20°C freezer for eight weeks and then returned to the greenhouse, and; (iv) three were fertilized again as above, and watered twice a week to potentially re-sprout. A reference specimen was collected in May 2022 from experiment (iv) and is deposited in the Purdue University Kriebel Herbarium.

## RESULTS

By May 2022 there was no evidence that the plants placed outside had survived the winter (Fig 1C). Average monthly temperatures recorded at the Purdue Agronomy Center for Research and Education (ACRE, located 8 km from the greenhouse; 40°47’00.3” N, 86°99’19.2” W) from November 2021 to March 2022 were 4.3°C, 4.2°C, −5.2°C, −2.2°C and 5.4°C respectively, and the minimum daily temperature recorded during this period was −9.8°C.

Following removal from the −20°C freezer after either four or eight weeks, plants were returned to the greenhouse and watered for 8 weeks to observe growth. here was no evidence that any plants had survived either four or eight weeks at −20°C (Fig 1D).

The plants that were kept in the greenhouse and never exposed to frost re-sprouted from clipped stems (Fig 1E) and by May 2022 still showed no signs of senescence (Fig 1F). Careful observation was made as to the source of the new vegetative growth, and they clearly had emerged from the cut stems and not as newly germinated seeds (Fig 1E). I note that plants flowered for the first time in September 2021, when the photoperiod was approximately 12-13 hours at this latitude. The photoperiod at the time of the second flowering was also approximately 12-13 hours, this time in April 2022. Thus, at the time of this writing those plants were 15 months old, had reproduced twice, and showed no signs of senescence.

## DISCUSSION

The results of the experiments described above lead me to conclude that *E. capillaris* is only an annual plant in those years or locations where it is killed by frost. Based on these results I suggest that the duration of *E. capillaris* should be revised to be “annual or perennial” in identification guides to match similar species that can sometimes exhibit an annual duration, and sometimes exhibit a perennial duration.

This new natural history information has potential implications for the study of the ecology and evolution of *E. capillaris.* The native range of *E. capillaris* includes all of eastern north America from Texas, to Florida USA in the south, up into Ontario and Quebec, Canada in the north, and as far west as Montana, USA (Bourfford 1993+; Usda & Nrcs 2022). The lack of frost tolerance supports the observation that *E. capillaris* grows as an annual throughout much of this range. However, my observations show that *E. capillaris* has the potential to survive and reproduce multiple times in the absence of a killing frost. This means it has the potential to grow more like a perennial in the southern part of its range. Additionally, with a warming climate I hypothesize that *E. capillaris* could begin to exhibit a perennial duration in more locations and years.

Those interested in the evolution of perennation or frost tolerance might examine *E. capillaris* as a potential model species for such work. I note that the seeds used in this study were sourced from Kentucky, USA, which is near the southern edge of the range of *E. capillaris.*A study of genetic variation across its range could also prove interesting to examine further evolutionary insights into the genetic basis of perennation or frost tolerance. Indeed, a warming climate is likely to lead to large changes in the ecology of many species that can be annual or perennial and so it may turn out that more species which were historically annual in duration might transition to perennial duration.

## ACKNOWLEDGEMENTS

I thank Scott McAdam and Chris Oakley for advice on these experiments. This work was funded by USDA National Institute of Food and Agriculture Hatch Project 1010722.

## REFERENCES

Beatley, J.C. (1970) Perennation in astragalus lentiginosus and tridens pulchellus in relation to rainfall. Madro&#xfl;o, 20, 326–332.

Bourfford, D. (1993+) Eragrostis capillaris. Flora of north america north of mexico (ed. Flora-of-North-America-Editorial-Committee), New York and Oxford, vol. 25, pp. 79.

Fox, G.A. (1990) Perennation and the persistence of annual life histories. The American Naturalist, 135, 829–840.

Friedman, J. & Rubin, M.J. (2015) All in good time: Understanding annual and perennial strategies in plants. American Journal of Botany, 102, 497–499.

Koornneef, M. & Meinke, D. (2010) The development of arabidopsis as a model plant. The Plant Journal, 61, 909–921.

Sano, Y., Morishima, H. & Oka, H.-I. (1980) Intermediate perennial-annual populations oforyza perennis found in thailand and their evolutionary significance. The botanical magazine = Shokubutsu-gaku-zasshi, 93, 291–305.

Usda & Nrcs (2022) The plants database (http://plants.Usda.Gov, 05/21/2022). National plant data team, greensboro, nc USA.

